# Adrenergic-melatonin heteroreceptor complexes are key in controlling ion homeostasis and intraocular eye pressure and their disruption contributes to hypertensive glaucoma

**DOI:** 10.1101/636688

**Authors:** Hanan Awad Alkozi, Gemma Navarro, David Aguinaga, Irene Reyes-Resina, Juan Sanchez-Naves, Maria J Perez de Lara, Rafael Franco, Jesus Pintor

## Abstract

Melatonin regulates intraocular pressure (IOP) whose increase leads to glaucoma and eye nerve degeneration. Aiming at elucidating the role of melatonin receptors in humour production and IOP maintenance, we here demonstrate that glaucoma correlates with disassembly of α_1_-adrenergic/melatonin receptor functional units in cells producing the aqueous humour. Remarkably, α_1_-adrenoceptor-containing complexes do not coupled to the cognate Gq protein and, hence, phenylephrine activation of these receptors does not lead to Ca^2+^ mobilization. Functional complexes are significantly decreased in models of glaucoma and, more importantly, in human samples of glaucoma patients (GP). In such glaucomatous conditions phenylephrine produces, via α_1_-adrenoceptor activation, an increase in cytoplasmic [Ca^2+^] that is detrimental in glaucoma. The results led to hypothesize that using melatonin, a hypotensive agent, plus blockade of α_1_-adrenergic receptors may normalize pressure in glaucoma. Remarkably, co-instillation of melatonin and prazosin, a α_1_-adrenergic receptor antagonist, results in long-term decreases in IOP in a well-established animal model of glaucoma. The findings are instrumental to understand the physiological function of melatonin in the eye and its potential to address eye pathologies by targeting melatonin receptors and their complexes.

## 1. Introduction

Glaucoma, a pathology characterized by visual field loss, is associated with optic nerve damage (Casson, Chidlow, Wood, Crowston, & Goldberg, 2012). The main risk factors are *inter alia* aging, genetic conditions and intraocular pressure (IOP) (Quigley, 2011). Besides being the second cause of blindness in the World, 61 million people suffer from glaucoma and by 2020 the number may approach 80 million (Quigley & Broman, 2006).

Normotensive IOP in adults is approximately 16 mmHg. Ocular hypertension is diagnosed when the value exceeds 21 mmHg (Li, Huang, & Zhang, 2018). Persistent ocular hypertension results in damage of the optic disc, causing degeneration of ganglion cells. The relationship between ocular hypertension and glaucomatous pathology is a well-known phenomenon (Sihota, Angmo, Ramaswamy, & Dada, 2018). However, despite the current therapeutic arsenal to decrease IOP, ocular hypertension is still the most important risk factor for optic nerve degeneration (Rossetti et al., 2015; Tamm, Braunger, & Fuchshofer, 2015).

Parasympathomimetics, adrenergic receptor antagonists, carbonic anhydrase inhibitors and prostaglandins reduce ocular hypertension (Hommer, 2010; D. A. Lee & Higginbotham, 2005) although may present notorious side effects (Beckers, Schouten, Webers, van der Valk, & Hendrikse, 2008). Better and safer interventions include drug co-administration as a way to reduce doses and side effects (Polo, Larrosa, Gomez, Pablo, & Honrubia, 2001; Sakai et al., 2005).

Anti-glaucomatous compounds reduce aqueous humour formation by the ciliary body thus leading to decreased IOP. In the healthy eye, the control of aqueous humour production is tightly controlled by hormones/neuromodulators, and aging leads to imbalance and increased hydrostatic pressure (Delamere, 2005). The role of adrenoceptors expressed in the ciliary body seems to be opposite as either α-adrenergic agonists or β-adrenergic antagonists may decrease IOP (Kiuchi, Yoshitomi, & Gregory, 1992; Murray & Leopold, 1985; Naito, Izumi, Karita, & Tamai, 2001). Studies in different animal models indicated that epinephrine, i.e. the endogenous agonist, produces a reduction in IOP, which can be blocked by prazosin, a selective α_1_-adrenoceptor antagonist (Funk, Wagner, & Rohen, 1992; P. F. Lee, 1958; Moroi, Hao, Inoue-Matsuhisa, Pozdnyakov, & Sitaramayya, 2000). Interest in melatonin is emerging in ocular diseases as it modulates aqueous humour production in the ciliary body. Thus, this indoleamine has potential in the treatment of ocular hypertension likely by acting via specific receptors: MT_1_ and MT_2_ (Crooke, Colligris, & Pintor, 2012; Mediero, Alarma-Estrany, & Pintor, 2009). The aim of this paper was to decipher in the eye the interplay of melatonin and epinephrine.

## 2. Material and methods

### 2.1 Human eye postmortem samples

Three eyes of normal subjects and two of glaucoma patients (GP) came from donations managed by the Balearic Islands tissue bank Foundation. After fixation with 4% paraformaldehyde, frontal sections (10 μm thick) were collected and stored at −20 ° until use.

### 2.2 Animals and intraocular pressure measurements

Experiments were performed using female C57BL/6J (n=5) (control) and DBA/2J (n=5) (glaucoma model) mice delivered by Charles River Lab. Institutional and regional Ethic Committees approved all procedures that included ARVO Statement for Ophthalmic and Vision Research. Mice were anesthetized by inhalation of isoflurane and IOP determined as previously described (Martinez-Aguila, Fonseca, Perez de Lara, & Pintor, 2016).

### 2.3 Cells, fusion proteins and expression vectors

The human non-pigmented ciliary epithelial cell line, 59HCE, was kindly supplied by Dr. Coca-Prados (Yale University) and grown in 10% high-glucose Dulbecco’s-modified Eagle’s medium (Gibco/Invitrogen, Carlsbad, CA). HEK-293T cells were grown in 5% FBS DMEM (Gibco/Invitrogen, Carlsbad, CA) as previously described (Martinez-Pinilla et al., 2017; Navarro et al., 2018).

The human cDNAs for the MT_1_, MT_2_ and a_1A_ receptors cloned in pcDNA3.1 were amplified without their stop codons using sense and antisense primers harboring either unique Hind III and BamH1 sites (MT_1_, MT_2_) or EcoRI and BamHI sites (α_1A_). The fragments were then subcloned to be in frame with a peYFP (YFP) and a pGFP^2^ (GFP^2^) placed on the C-terminal end of the receptor.

### 2.4 Transient transfection and sample preparation for energy-transfer experiments

HEK-293T cells growing in 6-well dishes were transiently transfected with the cDNA encoding for each protein/fusion proteins. Cells were incubated in serum-free medium (4h) with the corresponding cDNA, with polyethylenimine (PEI, 5.47 mM in nitrogen residues) and 150 mM NaCl. The rest of procedure was performed as described elsewhere for FRET, BRET and fluorescence determinations (Hinz et al., 2018; Martinez-Pinilla et al., 2017; Navarro et al., 2018).

### 2.5 cAMP, calcium determination, β-arrestin recruitment and label-free dynamic mass redistribution (DMR) assays

For cAMP determination 59HCE cells or HEK-293T cells were detached and re-suspended in medium containing 50 μM zardaverine (2,500 cells/well), pretreated (15 min) with the indicated molecule(s) and stimulated with agonists (15 min) before addition of 0.5 μM forskolin or vehicle. For [Ca^2+^] determination cells were transfected with the cDNA encoding for the GCaMP6 calcium sensor (3 μg cDNA) using lipofectamine 2000 (Thermo Fisher Scientific). 48 hours after transfection, cells (150,000 cells/well in 96-well black, clear bottom plates) were incubated with Mg^2+^-free Locke’s buffer pH 7.4 supplemented with 10 μM glycine and receptor ligands were added as indicated. To determine arrestin recruitment, BRET experiments were performed in HEK-293T cells 48 h after transfection with the cDNA corresponding to β-arrestin-2-Rluc (1 μg) and α_1A_-YFP (1,5 μg), or to β-arrestin-2-Rluc (1 μg) and MT_1_-YFP (1,5 μg), or to MT_2_-YFP (1,5 μg) alone or with a_1A_ (0.05 to 0,5 μg). Further details on procedures are elsewhere described (Martinez-Pinilla et al., 2017; Navarro et al., 2018).

### 2.6 Immunofluorescence and in situ proximity ligation (PLA) assays

Frozen sections from healthy and glaucoma subjects were rinsed in phosphate buffer saline (PBS) 1X and permeabilized with PBS-0.05% Tx-100 solution for 30 min. After blocking, antibodies raised against MT_1_ (1:200, Santa Cruz sc-13179), MT_2_ (1:1,000, ABIN122307, antibodies-online) and α_1_-adrenergic (1:500, Abcam ab3462) receptors were used. The rest of the protocol was similar to that elsewhere described using *ad hoc* secondary regents. PLA allows the *ex vivo* detection of molecular interactions between two endogenous proteins. PLA probes were obtained by linkage of primary anti-MT_1_ or MT_2_ receptor antibodies to PLUS oligonucleotide (DUO92009, Sigma, St. Louis, USA) and the α_1A_-adrenergic antibody to MINUS oligonucleotide (DUO92010, Sigma, St. Louis, USA). Samples were analyzed using confocal microscope (Zeiss LSM 5, Jena, Germany) at 40X magnification. The rest of the protocol was performed as described elsewhere, red spots were counted in each of the ROIs obtained in the nuclei images and data analysis was performed using specific PLA software (Franco et al., 2018; Martinez-Pinilla et al., 2017; Navarro et al., 2018).

## 3. RESULTS

### 3.1 Melatonin and α_1A_-adrenergic receptors in 59HCE cells are uncoupled from their cognate G proteins

Melatonin acts via MT_1_ (MT_1_R) and MT_2_ (MT_2_R) receptors, which control Cl^-^ efflux from ciliary body epithelial cells (Huete-Toral, Crooke, Martinez-Aguila, & Pintor, 2015). Interestingly, prazosin blocks the effect of melatonin on reducing IOP. As it is a α_1A_-adrenoceptor (α_1A_-AR) antagonist (Dubocovich, 1995; Huete-Toral et al., 2015; Pintor, Pelaez, Hoyle, & Peral, 2003) we hypothesized that melatonin and α_1A_-adrenergic receptors (α_1A_AR) could interact.

Cognate heterotrimeric G proteins are G_i_ for melatonin receptors, and G_q/11_ for α_1A_-AR (www.guidetopharmacology.org). We then measured, in 59HCE ciliary body epithelial cells, cAMP levels and dynamic mass redistribution (DMR). The effect of MT_1_ or MT_2_ receptor agonists did not result in decreasing forskolin-induced cytosolic [cAMP] (Fig. 1A-B). When cells were challenged with the α_1A_AR agonist phenylephrine, a remarkable elevation of cytosolic forskolin-induced cAMP was achieved. The antagonist prazosin abolished the cAMP elevation produced by phenylephrine. We also assayed the canonical coupling of α_1A_AR to heterotrimeric G_q_ proteins. We however did not find any effect on Ca^2+^ levels in 59HCE cells treated with phenylephrine (Fig. 1C). These results demonstrate that human ciliary epithelial cells express melatonin MT_1_ and MT_2_, and α_1A_-adrenergic receptors not coupled to cognate G proteins. Adrenoceptor mediated increases in cAMP reflect a Gs coupling able to overcome the effect of forskolin. It should be noted that the concentration of forskolin chosen for the assays (500 nM) is optimal as it allows detection of both Gi- and Gs-coupling.

**Figure 1.**
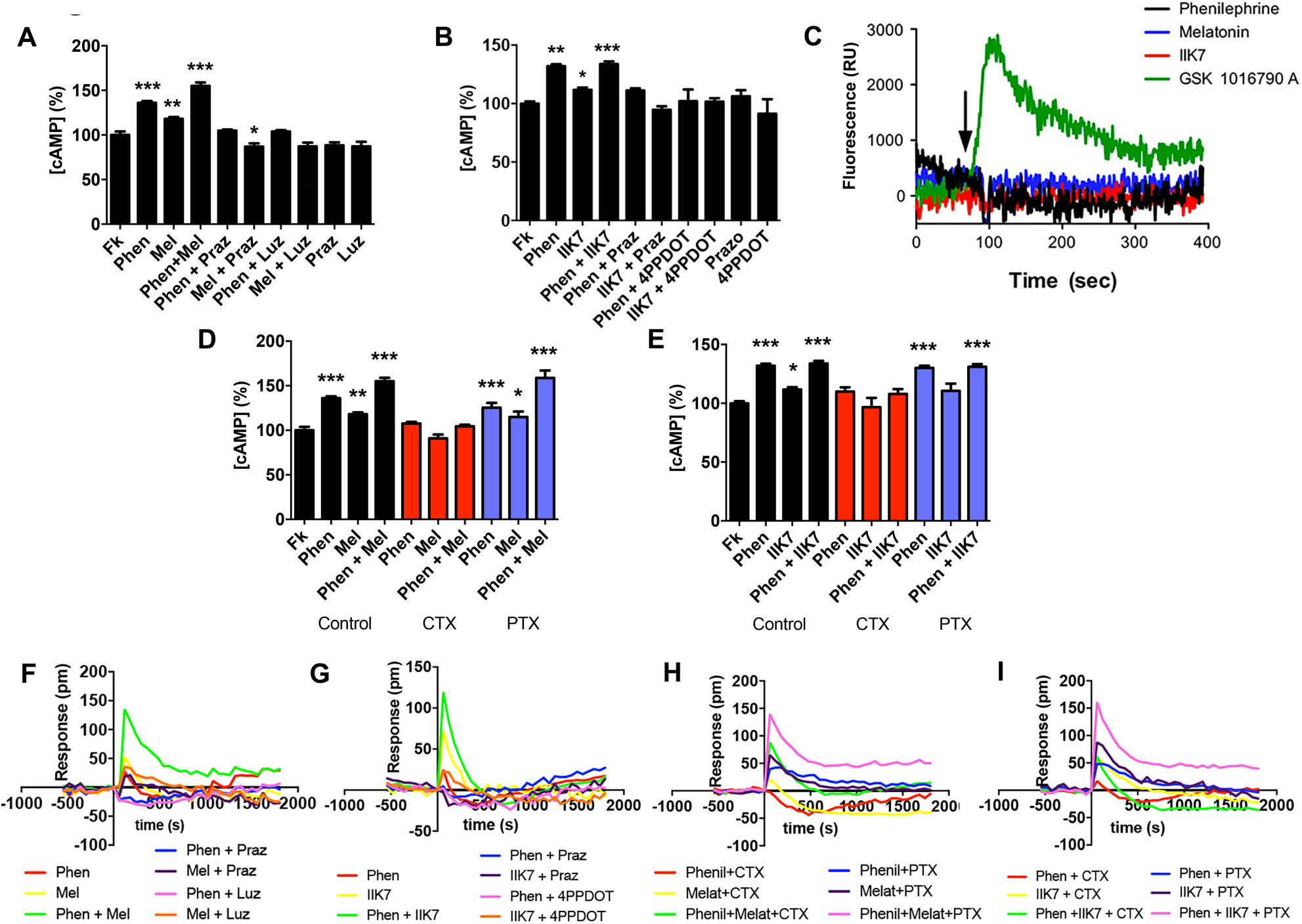
Effect of melatonin receptor agonists and of phenylephrine in human 59HCE cells. Panels A, B, D and E: Effect of ligands (single or combined treatment) on 0.5 μM forskolin-induced cAMP levels in the absence (A-B) or presence (D-E) of 10 ng/mL pertussis (overnight) or 100 ng/mL cholera (2 h) toxins. Data are given in percentage (100% represents the forskolin effect); they are the mean ± SEM (n=12, each in triplicates). One-way ANOVA followed by Bonferroni’s multiple comparison *post-hoc* test were used for statistical analysis. *p<0.05; **p<0.01; ***p<0.001. Panel C: Time course of cytosolic calcium levels induced by receptor agonists or by GSK-1016790A, an agonist of TRPV4 channels used for positive control. Panels F-I: Effect of ligands (single or combined treatment) on dynamic mass redistribution (DMR) in the absence (F-G) or presence (H-I) of pertussis or cholera toxins. Concentrations in the assays were: 100 nM phenylephrine, 1 μM melatonin, 100 nM IIK7, 1 μM prazosin, 1 μM luzindole and 1 μM 4PPDOT.

To confirm atypical G protein coupling, similar assays were performed in the presence toxins that disrupt G_s_- or G_i_-mediated signaling. Results show that cholera but not pertussis toxin inhibited the action of agonists, unequivocally indicating G_s_ involvement (Fig. 1D, 1E). Interestingly, the presence of the α_1A_-adrenergic antagonist, prazosin, abolished any effect of melatonergic agonists. Further evidence of cross-antagonism, a well-accepted heteromer print (Franco, Martinez-Pinilla, Lanciego, & Navarro, 2016), was underscored using a label-free technique consisting of detecting cell dynamic mass redistribution (DMR). The signal induced by any agonist was relatively high, and the combination of phenylephrine and either melatonin (Fig. 1F) or IIK7 (Fig. 1G), provided a more robust DMR response. As in cAMP determination assays, a cross-antagonism between α_1A_AR and melatonin receptors was detected. When the experiments were carried out in the presence of cholera and pertussis toxins, responses were again abolished by cholera toxin (Fig. 1H-I). Taken together, these results suggest that the crosstalk between melatonergic and α_1A_-adrenergic receptors are solely due to coupling to G_s_.

### 3.2 Atypical signaling in ciliary cells is due to α_1A_-adrenergic/melatonin receptor complexes

To identify potential direct interactions, a FRET biophysical approach was used (Fig. 2A). A saturable FRET curve was obtained in HEK-293T cells transfected with constant [cDNA] for α_1A_-ARGFP^2^ and increasing [cDNA] for MT_1_R-YFP (FRET_max_ 49 mFU and FRET_50_ 44) (Fig. 2B). Similar experiments using increasing amounts of [cDNA] of MT_2_R-YFP also provided a saturable FRET curve with FRET_max_ and FRET_50_ values of, respectively, 168 mFU and 128 (Fig. 2C). Thus, both melatonin receptors may form heteromers with α_1A_AR (α_1A_-MT_1_Hets and α_1A_-MT_2_Hets).

**Figure 2.**
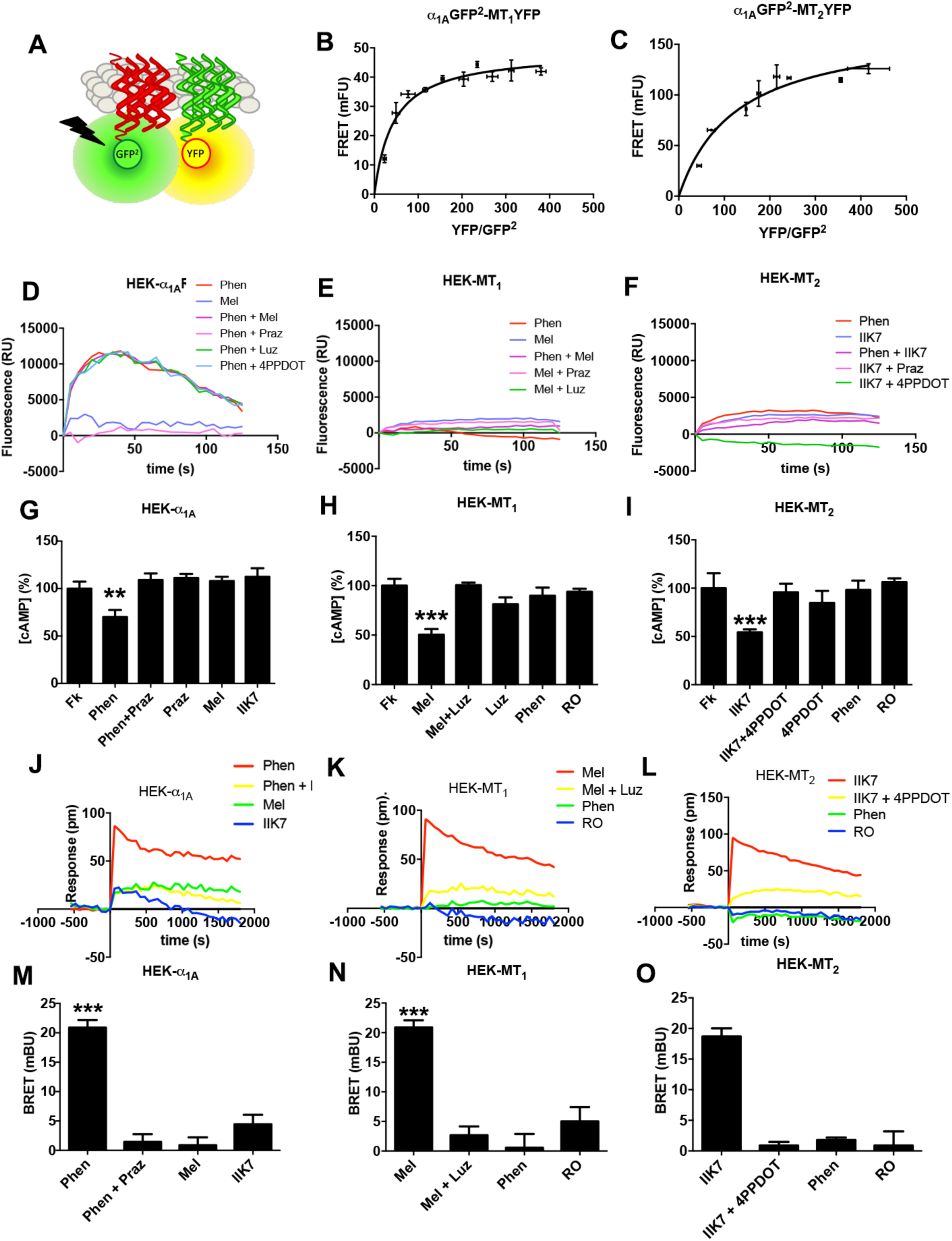
Dimerization of α_1_-adrenergic and MT_1_ or MT_2_ receptors and signaling via adrenergic or melatonin receptors in a heterologous expression system. Panels A-C: Scheme of the FRET assays (A) and results of experiments performed in HEK-293T cells transfected with a fixed amount of cDNA for the α_1_-adrenergic-GFP^2^ fusion protein and increasing amounts of cDNA for either MT_1_-YFP (B) or MT_2_-YFP (C). Energy transfer data are given in milliFRET units (mFU). The remaining experiments were performed in HEK-293T cells transfected with cDNAs for α_1_-adrenergic, MT_1_ or MT_2_ receptors. Panels D-F: Time course of cytosolic calcium levels induced by receptor agonists (single or combined treatment; when indicated, a preincubation with antagonists was performed). Panels G-I: Effect of receptor ligands (single or combined treatment) on 0.5 μM forskolin-induced cAMP levels. Data are given in percentage (100% represents the forskolin effect); they are the mean ± SEM (n=12, each in triplicates). For negative control, a selective agonist for dopaminergic D_4_ receptor, RO-105824, was used. One-way ANOVA followed by Bonferroni’s multiple comparison *post-hoc* test were used for statistical analysis. **p<0.01; ***p<0.001. Panels J-L: Effect of ligands (single or combined treatment) on dynamic mass redistribution (DMR). Panels M-O: BRET-based measurements of the effect of receptor agonists (single or combined treatment) on ß-arrestin 2-Rluc recruitment to every receptor-YFP fusion protein. Data are given in milliBRET Units (mBU); they are the mean ± SEM (n=10, each in triplicates). One-way ANOVA followed by Bonferroni’s multiple comparison *post-hoc* test were used for statistical analysis. ***p<0.001. Concentrations in the assays in panels D to O were: 100 nM phenylephrine, 1 μM melatonin, 100 nM IIK7, 1 μM prazosin, 1 μM luzindole, 1 μM 4PPDOT and 100 nM RO-105824.

Receptor functionality was first addressed in single transfected cells. Upon α_1A_AR activation a robust increase in cytosolic [Ca^2+^] detected by the fluorescence due to calcium-bound to an engineered GCaMP6 calmodulin sensor was obtained (Fig. 2D). The agonist effect was inhibited by prazosin, but not by the melatonin receptor antagonists, luzindole or 4-PPDOT. Neither MT_1_ nor MT_2_ receptors were coupled to G_q/11_ in single-transfected cells (Fig. 2E-F). Experiments of determination of forskolin-induced cAMP levels showed a G_i_-protein-coupling (Fig. 2G-I). The results confirm that, when expressed individually, receptors couple to their cognate G proteins. Activation of receptors in single transfected cells also provided significant DMR read-outs (Fig. 2J-L). Finally, BRET assays performed using β-arrestin-2-Rluc and receptors fused to YFP showed recruitment to either α_1A_, MT_1_ or MT_2_ receptors when selective agonists were used (Fig. 2M-O).

G protein coupling was completely different in co-transfected cells (Fig. 3A-B). On the one hand, phenylephrine did not produce Ca^2+^ responses in cells expressing either α_1A_-MT_1_Hets or α_1A_-MT_2_Hets (Fig. 3C-D). cAMP determination data in co-transfected cells were similar to the results obtained in 59HCE cells; agonists did not decrease forskolin-induced cAMP levels in cells expressing α_1A_-MT_1_Hets or α_1A_-MT_2_Hets. Also, cross-antagonism reappeared in cotransfected cells. Analogies between 59HCE and cotransfected HEK-293T cells were further found in experiments with toxins; the combined effect of agonists was blocked by cholera but not by pertussis toxin (Fig. 3E-F). Also, matching results in 59HCE cells, a cross-antagonism and a potentiation of the label-free DMR signal were found upon co-activation, and cholera toxin did block the effect (Fig. 3G-J). These results indicate a lack of productive coupling of receptor heteromers with G_q/11_ or G_i_. Cross-modulation was found in β-arrestin-2 recruitment but with a particular feature, namely both α_1A_-receptor activation and activation of MT_1_ melatonin receptors recruited β-arrestin to the α_1A_-receptor, co-activation resulting in a stronger signal (Fig. 3K). Antagonists of the two receptors abolished recruitment induced by agonists (cross-antagonism was, therefore, found). Similar results were obtained in cells co-expressing α_1A_-MT_2_ heteroreceptor complexes using *ad hoc* agonists/antagonists (Fig. 3L).

**Figure 3.**
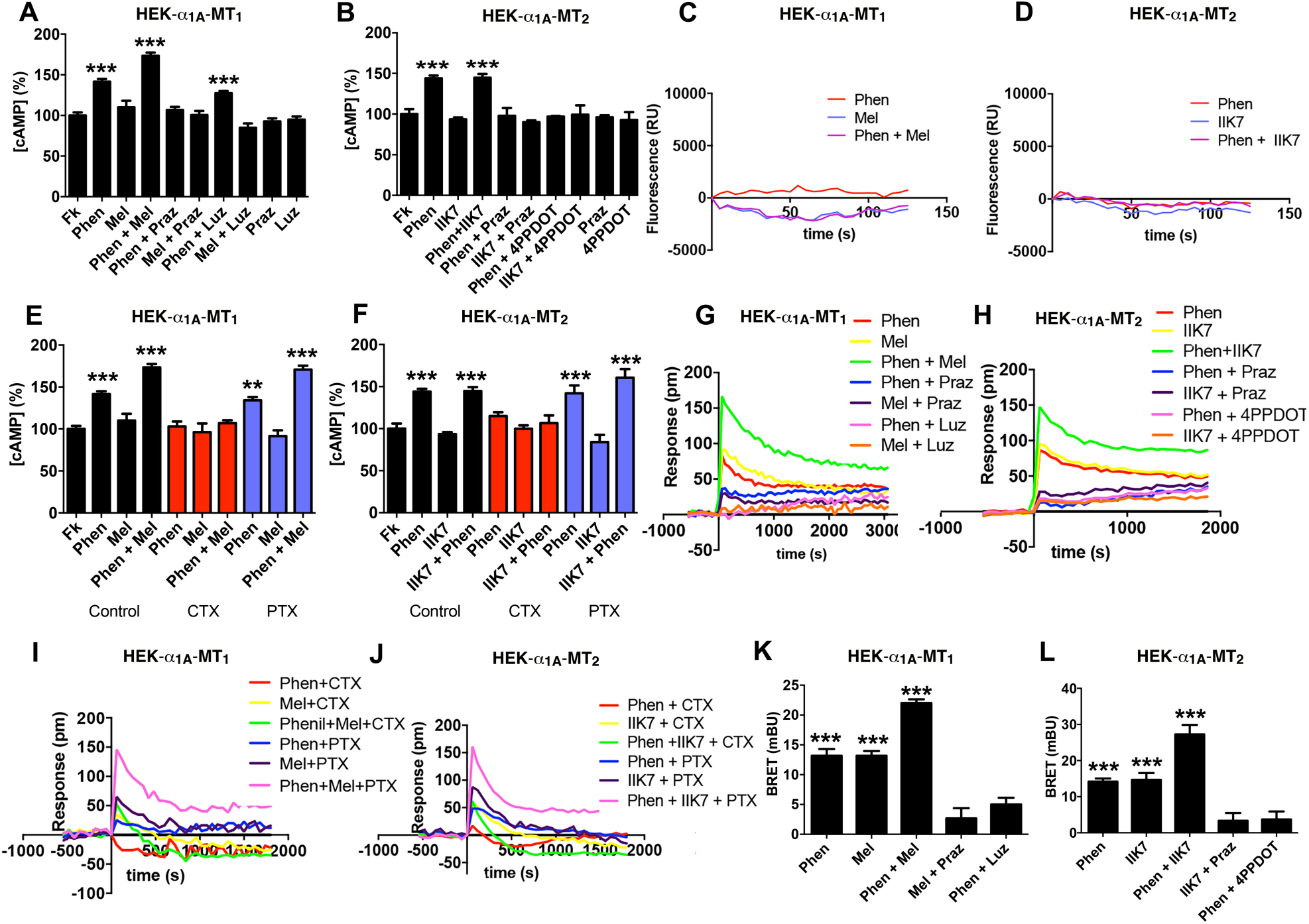
Signaling via adrenergic/melatonin receptor functional units in an heterologous expression system. Experiments were performed in HEK-293T cells transfected with cDNAs for full-length α_1A_-adrenergic and MT_1_ or MT_2_ receptors. Panels A, B, E and F: Effect of ligands (single or combined treatment) on 0.5 μM forskolin-induced cAMP levels in the absence (A-B) or presence (E-F) of 10 ng/mL pertussis (overnight) or 100 ng/mL cholera (2 h) toxins. Data are given in percentage (100% represents the forskolin effect); they are the mean ± SEM (n=12, each in triplicates). One-way ANOVA followed by Bonferroni’s multiple comparison *post-hoc* test were used for statistical analysis. **p<0.01; ***p<0.001. Panels C-D: Lack of effect of receptor agonist in cytosolic calcium levels. The agonist of TRPV4 channel, GSK-1016790A, did produce effect (data not shown). Panels G-J: Effect of ligands (single or combined treatment) on dynamic mass redistribution (DMR) in the absence (G-H) or presence (I-J) of pertussis or cholera toxins. Panels K-L: Effect of receptor agonists (single or combined treatment) on ß-arrestin 2 recruitment to MT_1_-YFP (K) or to MT_2_-YFP (L). Data are given in mBU and are the mean ± SEM (n=10, each in triplicates). One-way ANOVA followed by Bonferroni’s multiple comparison *post-hoc* test were used for statistical analysis. ***p<0.001. Concentrations in the assays were: 100 nM phenylephrine, 1 μM melatonin, 100 nM IIK7, 1 μM prazosin, 1 μM luzindole and 1 μM 4PPDOT.

### 3.3 The glaucomatous eye has reduced heteroreceptor complex expression and altered signaling

First of all, identification of α_1A_-MT_1_Hets and α_1A_-MT_2_Hets was achieved in healthy and glaucomatous conditions. The *in situ* proximity ligation (PLA) technique is instrumental to detect receptor-receptor interactions in a native system. Red clusters coming from PLA assays proved the occurrence of both α_1A_-AR/MT_1_R and α_1A_-AR/MT_2_R complexes in 59HCE cells. The analysis of the PLA labeling provided values of 65±10 dots/nucleus in the case of the α_1A_-MT_1_Hets and 73±8 dots/nucleus in the case of the MT_2_/α_1_ heteromer. The percentage of cells that presented positive PLA was 56±4 for α_1A_-MT_1_Hets and 57±5 for α_1A_-MT_2_Hets (n=150).

Does eye hypertension correlate with altered expression of melatonin-adrenoceptor complexes? We approached this question using cells subjected to stimulation of the transient receptor potential vanilloid 4 (TRPV4) channel. As previously reported (H. A. Alkozi, Perez de Lara, Sanchez-Naves, & Pintor, 2017; H. A. Alkozi & Pintor, 2015), activation of the channel mimics the ion fluxes that drive the increase in hydrostatic pressure that occurs in the hypertensive/glaucomatous eye. The application to 59HCE cells of the TRPV4 agonist, GSK1016790A, modified the PLA signal in a dose-dependent manner. As shown in Fig. 4A-B, the higher the concentration of GSK1016790A, the lower the PLA signal. Therefore, a reduction in the expression α_1A_-MT_1_Hets or α_1A_-MT_2_Hets happens when a glaucomatous condition is reproduced in a preclinical model (Fig. 4C).

**Figure 4.**
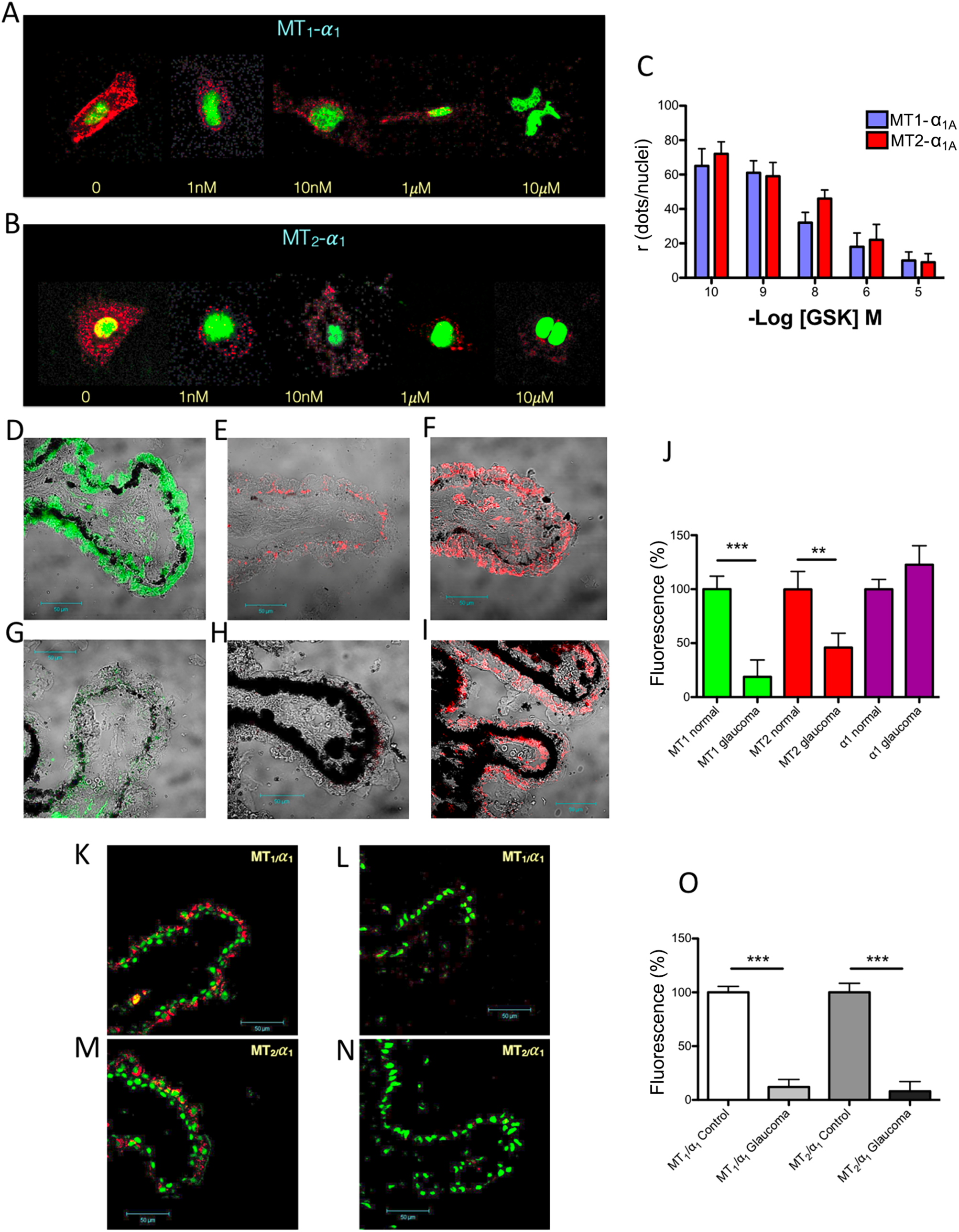
Identification of functional units in human 59HCE cells and in samples from normotensive and hypertensive human eyes. Panels A-B: Determination by PLA of α_1_-adrenergic and either MT_1_ (A) or MT_2_ (B) receptor complexes in 59HCE non-pigmented ciliary body epithelial cells. Panel C: Quantification of clusters for α_1_-adrenergic and either MT_1_ (blue) or MT_2_ (red) receptors. Data are the mean ± SEM (n=5). *p<0.005, using Student’s t-test. Values for negative control were, respectively, 5±1 and 2±1 dots/nucleus). Panels D-I. Immunolocalization of MT_1_ (D), MT_2_ (E) or α_1_-adrenergic (F) receptors in the ciliary body of healthy controls. Immunolocalization of MT_1_ (G), MT_2_ (H) or α_1_-adrenergic (I) receptors in the ciliary body of GP. Quantitation (I) of results comparing patients and controls using fluorescence values taken in identical experimental conditions. Data, given in percentage (100% given to values from controls) are the mean ± SEM (n=4). **p<0.01 and ***p<0.001, using Student’s t-test. Panels K-O: Determination by PLA of α_1_-adrenergic and either MT_1_ (K) of MT_2_ (M) receptor complexes in the ciliary body of age-matched IOP normotensive individuals. Determination by PLA of α_1_-adrenergic and either MT_1_ (L) of MT_2_ (N) receptor complexes in the ciliary body of GP. Quantitation (O) of clusters comparing data in GP and controls using fluorescence values taken in identical experimental conditions. Data, given in percentage (100% given to values from controls) are the mean ± SEM, n=4 (***p<0.001, using Student’s t-test).

Heteromer expression was assayed in unique postmortem samples obtained from GP and age-matched controls. In the eye from healthy donors, the presence of melatonin MT_1_, MT_2_ and α_1A_-adrenergic receptors was confirmed by immunoreactivity across the ciliary body (Fig. 4D-F). A strong labeling for the MT_1_R was present in non-pigmented epithelial cells, while the labeling for the MT_2_R was observed in the basal membrane of the non-pigmented epithelium (Fig. 4D-E). Melatonin receptors were not found in the stroma. Concerning the α_1A_-adrenergic receptor, a positive labeling was observed in the pigmented and non-pigmented epithelial cells as well as in the stroma (Fig. 4F). Similar immunohistochemical studies carried out in samples obtained from GP showed altered receptor expression (Fig. 4G-I). The expression of the α_1_-adrenergic receptor showed a trend (to increase) that was not statistically significant (Fig. 4J). In sharp contrast, immunoreactivity for melatonin receptors was markedly reduced, 81±15 and 54±13% for, respectively, MT_1_ and MT_2_ receptors. For MT_1_/α_1A_ and MT_2_/α_1A_ receptor pairs, a marked PLA positive labeling was observed in control samples (Fig. 4K and 4M). Remarkably, when PLA was performed in samples from GP, the reduction of heteromers was 88% in the case of the MT_1_/α_1A_ and 90 % in the case of the MT_2_/α_1A_ heteromers (Fig. 4L, 4N and 4O). Once more, these results match with the results obtained in 59HCE cells treated with a TRPV4 activator to mimic a glaucoma-like condition.

### 3.4 A novel therapeutic approach to combat glaucoma

In GP eyes and in the eye of glaucomatous models, the coupling of α_1A_-adrenoceptors to G_q_ and the subsequent Ca^2+^ signaling seems detrimental and markedly contributing to the disease. Accordingly, an antagonist of α_1A_-AR would be beneficial as a blocker of calcium production and of Ca^2+^-regulated chloride channels (Fleischhauer, Mitchell, Peterson-Yantorno, Coca-Prados, & Civan, 2001). To test the hypothesis we moved to a well-established murine model of glaucoma (Perez de Lara et al., 2014). 3-month-old DBA/2J mice have normotensive eyes and physiological levels of melatonin. Retinal electrophysiology parameters were undistinguishable from those in the control mouse (C57BL/6 background) (Fig. 5A-C), however prazosin antagonized hypotensive effect of melatonin (Fig. 5D). 9-month-old mice display a full glaucoma-like pathology (Fig. 5A-C). At 9 months of age, and despite elevated levels of melatonin (Fig. 5B), the glaucomatous eye of DBA/2J mice was sensitive to the hypotensive effect of exogenously added melatonin, which reduced the IOP from 16.6±0.6 to 11.8±0.4 mmHg. Remarkably, prazosin, enhanced melatonin hypotensive action (to 9.0 ± 0.5 mmHg) instead of antagonizing it (Fig. 5F). Moreover, the effect of prazosin lasted more than 6 hours, thus indicating a very appropriate therapeutic time window. These results open a new and easy-to-implement anti-glaucoma treatment, consisting of melatonin/prazosin co-administration, with few expectable side effects.

**Figure 5.**
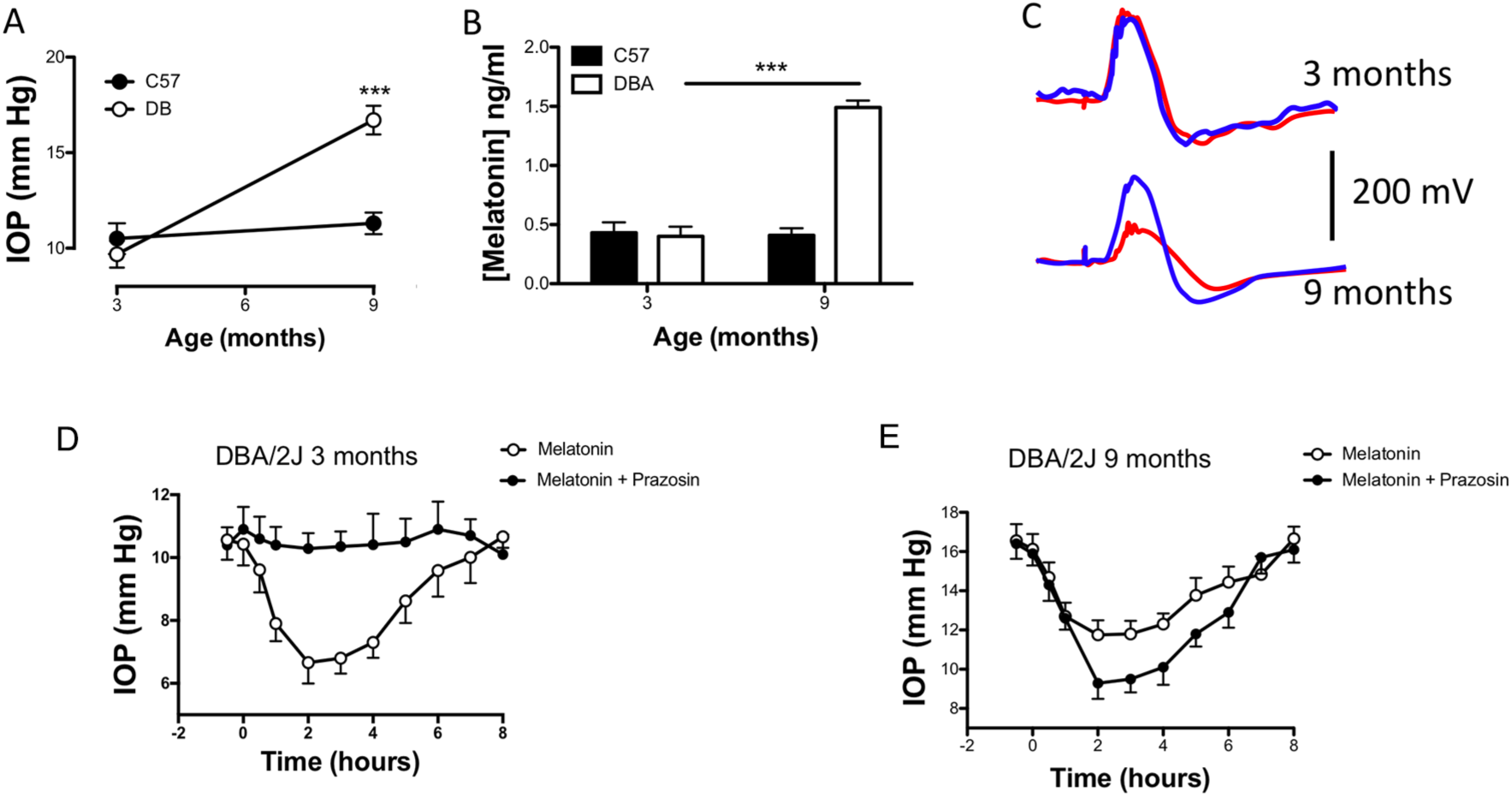
IOP-lowering intervention in the DBA/2J mice. Panel A: Intraocular pressure (IOP) values of control C57BL/6J and DBA/2J mice at 3 and 9 months of age. Data are the mean ± SEM (n=12). ***p<0.001 (Wilcoxon’s test for paired samples). Panel B: Melatonin concentrations in the aqueous humour of C57BL/6J and DBA/2J mice at 3 and 9 months of age. Data are the mean ± SEM (n=6). ***p<0.001 (Wilcoxon’s test for paired samples). Panel C: Electroretinogram in C57BL/6J (blue) and DBA/2J (red) mice. Positive scotopic threshold response (pSTR) amplitude was significantly reduced between 3 and 9 months of age (p<0.00001). Panels D-E: Time course of the effect on IOP in 3 (D) or 9 (E) month old DBA/2J mice after the instillation of melatonin ± prazosin. Data are the mean ± SEM (n=5). **p<0.01 (two-way ANOVA followed by *post-hoc* Tukey’s test).

## 4. Discussion

Current therapy of GP addresses symptoms by interventions neither efficacious in all patients nor absent of adverse events. Prostaglandin analogues have been approved for GP because they reduce intraocular pressure but by unknown mechanisms (Lanza et al., 2018; Sanford, 2014; Weinreb et al., 2015). Blockers of β-adrenoceptors or drugs able to engage α-adrenoceptors are in the portfolio; however, the rationale behind the hypotensive action of to ß-blockers and α-receptor agonists is unknown.

Melatonin, has become popular for potential properties for (among other) sleep induction and it is even on-line available. Already in 1993 (Serino, d’Istria, & Monteleone, 1993) described the production of melatonin in the pineal gland and in the retina by methylation of N-acetyl serotonin (Hardeland, 2010). Interestingly, melatonin decreases IOP but the significant increase in the aqueous humour of GP is not enough to normalize IOP (Alarma-Estrany, Crooke, Peral, & Pintor, 2007; H. Alkozi et al., 2017). Characterization of melatonin receptors has not been completed due to atypical pharmacological and non-fulfilled suspicions of a third receptor. We reasoned that atypical data could be due to interactions with other GPCRs as receptor heteromers display particular properties (Ferre et al., 2009). Here we selected α_1A_-AR because its activation leads to increases in cytosolic [Ca^2+^] that in turn control ion fluxes in the ciliary body. We here report that MT_1_ and MT_2_ receptors may directly interact with α_1A_-ARs.

The role of epinephrine in the physiology of the eye is known since 1970 (Drance & Ross, 1970; Zalta, Shock, Stone, & Petursson, 1983). Interestingly, the lack of coupling of receptors for epinephrine to G_q/11_ in the ciliary body (Figure 1), both intriguing and relevant, is due to α_1A_-Rs forming heteromers with melatonin receptors. Remarkably, the glaucomatous eye expresses few functional units and the signaling mediated by Gq-coupled α_1A_-ARs negatively impacts on IOP.

Our results predicted that the effect of melatonin would be enhanced by blockade of G_q_-coupled α_1A_-adrenergic receptors by prazosin. Indeed, combination of the two compounds led to normalize IOP in a well-established model of glaucoma and the effect was not transient but lasted for 6 hours. The long lasting experience with prazosin as blood pressure lowering agent indicates that it is very safe and does not display the serious side effects provoked by other α-adrenergic receptor antagonists (Brogden, Heel, Speight, & Avery, 1977).

In summary, we have discovered a trend in glaucoma consisting of the disruption of complexes formed by adrenergic and melatonin receptors. Such trend found in animal models of the disease and in samples from human eye (from GP and age-matched controls) makes ineffective the huge 3.2-fold increase in the concentration of melatonin in the glaucomatous eye. The physiological functional unit is coupled to a G_s_ protein whereas the disassembly leads to α_1A_-adrenergic receptors coupled to G_q_ and to melatonin receptors coupled to G_i_. Increases in calcium via G_q_ and decreases of cAMP via G_i_ establish a vicious circle that negatively impacts on the ion channels controlling intraocular pressure. The mechanism consisting of allosteric interaction and shift of G protein coupling is quite noteworthy and explains one early finding in the laboratory that is included in the paper, namely the lack of calcium ion mobilization by phenylephrine in cells expressing α_1A_-adrenoceptors (Fig. 1C). Together, our findings provide a better understanding of the ciliary body physiology, completing a preclinical translational research addressed to combat glaucoma. Remarkably, a therapeutic strategy resulting from combining melatonin, sold as a supplement and lacking collateral effects even at high doses in the eye (Rosenstein et al., 2010; Sanchez-Barcelo, Mediavilla, Tan, & Reiter, 2010), and prazosin, approved for the therapy of blood hypertension (Brogden et al., 1977; Mallorga, Buisson, & Sugrue, 1988; Singleton et al., 1989; Torvik & Madsbu, 1986), could readily enter into clinical trials to assay for safety and efficacy in humans.

## AUTHOR’S CONTRIBUTION

HAA and GN participated in the design and performance of many of the experiments and analyzed the results; it is considered that their contribution was similar. JSN was instrumental in obtaining and providing human samples for this study. MJPL performed IOP measurements. DA participated in immunohistochemistry and PLA assays. RF and JP designed and supervised the work; it is considered that their contribution was similar. HAA, GN, RF and JP actively participated in writing and editing. All authors have edited the paper and have received a copy of the final version.

## Acknowledgements

This work was supported by grants from Spanish *Ministerio de Economía y Competitividad* (MINECO) Ref. [SAF2013-44416-R], [SAF2016-77084-R] and [BFU2015-64405-R], and from Spanish *Ministerio de Sanidad* Ref. RETICS [RD12/0034/0003] and [RD16/ 0008/0017]. Hanan A. Alkozi is a fellowship holder of Saudi Arabia government. MINECO grants may include EU FEDER funds.

